# A novel multi-perspective imaging platform (M-PIP) for phenotyping soybean root crowns in the field increases throughput and separation ability of genotype root properties

**DOI:** 10.1101/309161

**Authors:** Anand Seethepalli, Larry M. York, Hussien Almtarfi, Felix B. Fritschi, Alina Zare

**Author notes:** Corresponding authors: Felix B. Fritschi, Alina Zare.

## Abstract

**Background:** Root crown phenotyping has linked root properties to shoot mass, nutrient uptake, and yield in the field, which increases the understanding of soil resource acquisition and presents opportunities for breeding. The original methods using manual measurements have been largely supplanted by image-based approaches. However, most image-based systems have been limited to one or two perspectives and rely on segmentation from grayscale images. An efficient high-throughput root crown phenotyping system is introduced that takes images from five perspectives simultaneously, constituting the Multi-Perspective Imaging Platform (M-PIP). A segmentation procedure using the Expectation-Maximization Gaussian Mixture Model (EM-GMM) algorithm was developed to distinguish plant root pixels from background pixels in color images and using hardware acceleration (CPU and GPU). Phenes were extracted using *MatLab* scripts. Placement of excavated root crowns for image acquisition was standardized and is ergonomic. The M-PIP was tested on 24 soybean [*Glycine max* (L.) Merr.] cultivars released between 1930 and 2005.

**Results:** Relative to previous reports of imaging throughput, this system provides greater throughput with sustained rates of 1.66 root crowns min^-1^. The EM-GMM segmentation algorithm with hardware acceleration was able to segment images in 10 s, faster than previous methods, and the output images were consistently better connected with less loss of fine detail. Image-based phenes had similar heritabilities as manual measures with the greatest effect sizes observed for Maximum Radius and Fine Radius Frequency. Correlations were also noted, especially among the manual Complexity score and phenes such as number of roots and Total Root Length. Averaging phenes across perspectives generally increased heritability, and no single perspective consistently performed better than others. Angle-based phenes, Fineness Index, Maximum Width, Holes, Solidity and Width-to-Depth Ratio were the most sensitive to perspective with decreased correlations among perspectives.

**Conclusion:** The substantial heritabilities measured for many phenes suggest that they are potentially useful for breeding. Multiple perspectives together often produced the greatest heritabilities, and no single perspective consistently performed better than others. Thus, as illustrated here for soybean, multiple perspectives may be beneficial for root crown phenotyping systems. This system can contribute to breeding efforts that incorporate under-utilized root phenotypes to increase food security and sustainability.

## Background

The global population is expected to increase to nine billion people by 2050 which necessitates an increase in global food production of at least 60%, but likely as much as 100% due to increased livestock production (Grafton et al. 2015). Soybean [*Glycine max* (L.) Merr.] is one of the major crops with a total global production of 351.32 million tons in 2016 (USDA 2018). Roots are crucial for plant productivity by foraging in soil for water and nutrients (Lynch 1995), yet have not been major targets of breeding efforts because a significant knowledge gap remains about specific relationships of root form and function with yield. Improving research capacity to measure root phenes, or elemental units of phenotype [1], could have major effects on breeding targets in soybean and other crops.

Advances in imaging and computing technologies have allowed efficient analysis of plant root images (Pound et al. 2013; Lobet et al. 2011). These methods use algorithms ranging from the more complex such as the EM-GMM algorithm (Dempster et al. 1977; Bilmes 1998) to simpler procedures to generate segmented images such as intensity thresholding (Colombi et al. 2015; Bucksch et al. 2014; Yugan and Xuecheng 2010; Huang et al. 1992). The segmented image may then be analyzed to extract root phenes. For example, Janusch et al. (2014) proposed to identify root topology using Reeb graphs, which depicts the topological structure of the root shape as the connectivity of level sets. Chen et al. (2006) described a method to determine the branching structure in wheat using Markov chains, where the lateral root branching probability was found using the locations of the lateral roots along the main root. However, these methods need relatively simple root systems to extract properties with accuracy. Mairhofer et al. (2012) and Zhou et al. (2014) described methods to perform 3D reconstructions from X-ray computed tomography by scanning the image stack vertically. Ying et al. (2011) performed reconstruction on RGB images using the regularized visual hull algorithm. Although complex root system architectures were successfully identified by the tool, the procedure is still computationally intensive and the image acquisition methods are not high-throughput. Plant roots grown in a gel medium were imaged while revolving on a turntable, yielding 3D phenotypes (Iyer-Pascuzzi et al. 2010; Clark et al. 2011). Pantalone et al. (1996) showed a relationship between drought tolerance and the level of complexity (fibrousness) of the root system. This work also tried to assign a root score to various root systems depending upon several factors such as the amount of fibrous root, and the size and number of root nodules. Topp et al. (2013) imaged rice root systems grown in clear gellan gum over a full rotation in order to reconstruct 3D models and extracted both 2D and 3D phenes.

Software exist that analyze root growth such as KineRoot (Basu et al. 2007) and ArchiSimple (Pagès et al. 2014) and using displacement vector fields (Kirchgessner et al. 2001). Balestri et al. (2015) discussed the changes in the root topology when a seagrass was grown in different soil conditions quantitatively. Further studies include modeling of roots (Jia et al. 2010), analysis of growth of roots in time series (Fang et al. 2009), imaging the roots by parts and stitching them (Kun et al. 2011) and estimation of root system architecture by modeling root length or number of roots per unit volume or root density (Dupuy et al. 2005).

Recently, two novel strategies were proposed for high-throughput phenotyping of plant roots, Digital Imaging of Root Traits (DIRT) (Bucksch et al. 2014) and Root Estimator for Shovelomics Traits (*REST*) (Colombi et al. 2015). The procedures consist of a root imaging standard methodology for excavated root crowns followed by image processing, phene extraction and analysis of the extracted phenes. The *DIRT* platform was meant to be robust enough to analyze images from various imaging methods and light conditions, while *REST* used an optimized imaging system using a blackout tent and flash lighting to produce easily segmentable images. These systems built on the recent innovations of manually scoring root crowns, sometimes called ‘shovelomics’ (Trachsel et al. 2011). In general botanical terminology, root crown refers to the site where the root system transitions to the shoot (Beentje 2010), and in the root phenotyping context root crown has generally been accepted to refer to both the crown itself and the entirety of attached roots following excavation. That is, the root crown is the top portion of the root system that remains after excavation and removal of soil by washing or other means. Root crown properties have been linked to crop performance, such as nodal root number (Saengwilai et al. 2014; York et al. 2013; York and Lynch 2015) and growth angle (Trachsel et al. 2013; York et al. 2015). The application of root crown phenotyping for legumes has recently been accomplished in common bean, soybean, and cowpea (Burridge et al. 2016a; Burridge et al. 2016b; Fenta et al. 2014). The use of these root crown phenotyping tools has led to the discovery of multiple phenes (fundamental units of phenotype, York *et al.*, 2013) that impact plant growth in the field.

The primary goal of this study was to create a high-throughput system that can extract image-based phenes from the photographed images of plant root crowns using custom hardware and software to optimize the throughput of image acquisition and analysis. Generally, root crowns have been imaged from a single perspective, however legume root crowns are much more asymmetric relative to cereal root crowns. Therefore, imaging from multiple perspectives may better reflect three dimensional aspects of soybean roots. Thus, a system combining a blackout box, internal lighting, image acquisition software, and five consumer cameras was developed and constitutes the Multi-Perspective Imaging Platform (M-PIP). In order to validate extracted phenes from the images, they were compared to manually measured or scored properties. Unlike phenotyping systems that take multiple images using a turntable (Iyer-Pascuzzi et al. 2010; Clark et al. 2011), the images were taken from multiple cameras in order to maximize throughput (no waiting for turntable revolution). Further, the new algorithm for segmentation advanced the state-of-the-art in root phenotyping by making use of hardware acceleration and working on color (RGB) images rather than grayscale. We show that with this combined setup, we can capture and process thousands of images per day.

## Methods

### Field experiment and root excavation

A field experiment including 24 maturity group IV soybean varieties released between 1930 and 2005 was conducted in 2016 in Columbia, MO, USA. The soil at the site is a Haymond silt loam (coursesilty, mixed, superactive, mesic Dystric Flueventic Eutrudepts) and the field was disked to approximately 0.15 m prior to planting. Soybean were planted at a density of 34 seeds m” on 7 May, 2016 in 3.05-m long four-row plots with 0.76 m distance between rows. The varieties were arranged in a randomized complete block design with four replications. Pre-emergence herbicide application and manual weeding were used to control weeds. At beginning seed (R5) (Fehr and Caviness 1977) five plants from one of the middle rows of each plot were cut 10 to 15 cm above the soil surface. To extract the roots from the soil, a circle with a radius of approximately 0.1 m centered on the stem was cut with a shovel and roots were excavated to a depth of 0.2 m. Soil attached to the roots was removed by shaking.

### Imaging protocol

The M-PIP was designed for acquiring images of the same plant root crown from different angles for phene extraction. The M-PIP consists of an imaging box designed for easy operability and transportability. The imaging box measures 137.2 cm (54 inch) wide, 101.6 cm (40 inch) tall, and 121.9 cm (48 inch) in depth and consists of a frame of T-slotted aluminum extrusion bars and three sides, a bottom and a top of thin plywood were attached to exclude light and block wind (Figure 1 A). The front side of the box was covered with black cloth that was moved aside to access the interior of the box with the cameras. An opening of 0.6 by 0.4 m was cut into the top of the imaging box and was used to insert the root crowns in the correct position for imaging. To this end, two lids were built to fit the opening in the top of the box and were fitted with a PVC pipe (1” and 0.34 m in length) mounted perpendicular to the plane of the lid and in the center of the lid. Plant stems were placed into the pipe and held in place with a piece of foam pipe insulation when needed, but often the curvature of the stem was sufficient to lock the root in place. This top-loading configuration of root crowns allowed ergonomic and rapid placement of root crowns in a defined location for repeatable imaging. The innovation further allowed high-throughput imaging by use of two lids, such that one root crown was being replaced while another was imaged.

**Figure 1.**
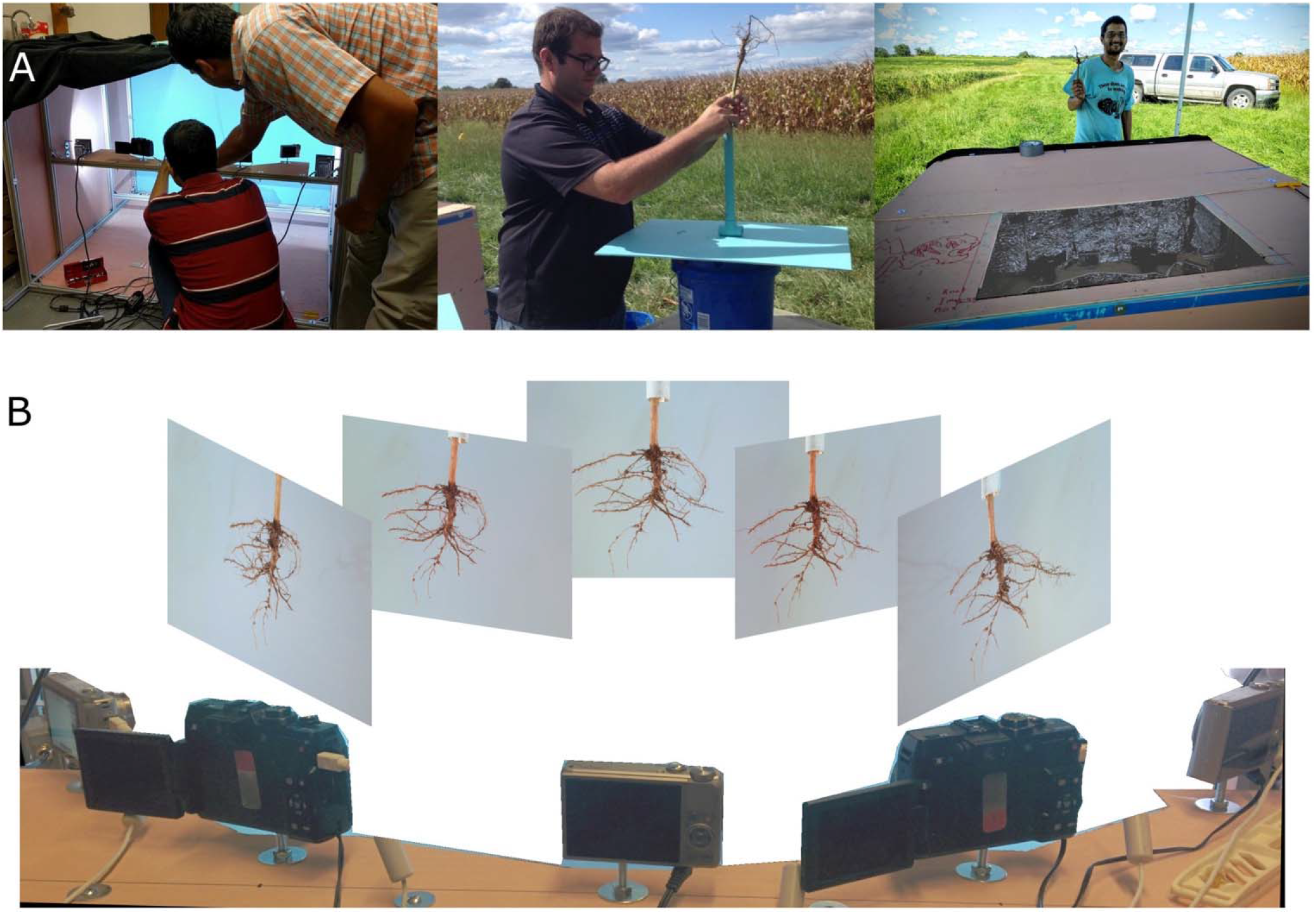
A. An imaging station was constructed from T-slotted aluminum extrusion and thin plywood. Internal lighting illuminated a suspended root crown against a blue-painted background. Root crowns were affixed to a PVC pipe painted the same as the background mounted to a top panel that could be placed over the hole in the back top surface of the imaging box in order to suspend the root crown in the focal position of the five cameras. B. Five cameras were placed at equal distances along a 90 degree arc. For clarity, the background has been deleted from the photo of the 5 cameras. The five root crown images are the five actual perspectives acquired by each camera.

The lid with the attached root crown was placed on the opening in the top of the imaging box, thus positioning the center of the root crown approximately 0.3 m from the plane of the background side. Images were taken from five cameras mounted on a wooden board inside the imaging box. The five cameras were placed along a circular arc equidistant (radius = 0.61 m) from the center of the plant root. The entire arc subtended an angle of 90° at the suspended plant root, and the cameras were placed at equal angles along this arc. When placing the roots in the M-PIP setup box, the perspective having the subjectively maximum spread was placed directly toward the central camera. This was to allow the cameras to capture the entire root structure, to keep consistency among root crowns, and to make sure the roots did not touch background. The cameras in the M-PIP system were configured to acquire images that optimized contrast of the roots with the blue background.

The background of the imaging box was painted with light blue to aid with image processing. Additionally, four daylight hued flood lamps (10W, Warmoon, Lightrace Technology Co., Ltd.) were installed to illuminate the root system suspended from the lid. To minimize shadows due to lighting, the illumination was adjusted to achieve diffuse light by the time it reached the background. Initially, aluminum foil was placed on the inside of the black curtain behind the cameras to reflect light and the cameras were placed closer to the background. As part of the optimization process aimed at facilitating image processing, the lights were moved farther away from the background, and the aluminum foil was replaced with a white shower curtain fabric.

The M-PIP system was powered by a generator in the field. The power was fed through AC to DC adaptors for each camera instead of batteries, which allowed continual use of the imaging system. All cameras were connected to a USB hub which was in turn connected to a PC. Canon Hacker Development Kit (CHDK) was installed on the SD cards inserted into the cameras. The CHDK was used to programmatically control the camera functions by sending commands from USB using Picture Transfer Protocol (PTP). The commands included changing the optical zoom, aperture, exposure time, ISO speed, capturing images and downloading the images to the PC over the USB connection. A client program was used to send commands to the Canon cameras from the PC. In this implementation for camera control, the cameras used were two Canon G1X cameras and three Canon S110 cameras. The Canon S110 cameras were placed at the middle and at either ends of the wooden board and the two G1X cameras were placed in between the S110 cameras, allowing visual comparison between the images at different angles because these images were obtained by cameras with the same image sensors. The Canon G1X cameras were operated at a focal length of 35 mm and the Canon S110 cameras at a focal length of 13.6 mm. The aperture was set to f/8.0 and the ISO speed was set at 800 for all cameras, and the exposure time was set to 1/40 seconds for Canon G1X cameras and 1/100 seconds for Canon S110 cameras. Once the images were taken, they were downloaded to the PC and segmentation was performed. In total, 480 (24 varieties x 4 replications x 5 roots per plot) soybean root crowns were imaged using the M-PIP. Given the five cameras used, 2400 images in total were acquired.

### Image segmentation

Image segmentation is needed to extract phenes from the plant root pixels located in the image. The images were first manually cropped to only background and the plant root pixels before performing segmentation. This involved cropping the clip or pipe used to hold the plant root at the time of imaging. The segmentation was performed using the Expectation Maximization - Gaussian Mixture Models (EM-GMM) algorithm (Dempster et al. 1977; Bishop 2006), by modeling the pixels as originating from two 3D Gaussian distributions. One Gaussian distribution models the background (blue color) pixels whereas the other distribution models the plant root pixels. The cropped image was passed to the EM-GMM algorithm to generate a segmented image. The largest connected component using flood-fill algorithm from the MATLAB Image Processing toolbox was selected from the segmented image to remove miss-classifications due to noise in the image and saved as the final segmented image.

The EM-GMM algorithm was implemented in C/C++ and accelerated through the use of NVIDIA Quadro K600 GPU with Compute Capability 3.0. Using our implementation on Intel E3-1271 v3 processor having quad-core CPU, which supports for Intel’s Advanced Vector Extensions 2.0 (AVX2) and Fused Multiply and Add (FMA) instruction sets, the program takes 25 seconds to segment a 4000 x 3000 pixel image. Whereas on NVIDIA Quadro K600 GPU, consisting of 192 CUDA cores, our GPU implementation takes only 10 seconds to segment the same image.

The multivariate EM-GMM algorithm was implemented such that the means of the two 3D Gaussian distributions were initialized from the peaks of the histograms of the red, green and blue channels of the image. Since, the images obtained from the cameras had a light blue background, the pixels that had a peak at a greater intensity in all the color channels were initially classified as background pixels. The remaining pixels were taken as the foreground or root pixels. Unlike most of the earlier works (Bucksch et al. 2014; Colombi et al. 2015) based on greyscale thresholding to separate root pixels from background, this EM algorithm auto-tunes the mean and the covariance parameters based on each image so that the likelihood of the pixels is maximized. Also, the algorithm segments an RGB image by estimating the full covariance matrix.

### Image-based Phene Extraction

Phenes were extracted from the segmented images to be used in statistical analysis later. The border pixels from the segmented image were identified and counted for perimeter. The segmented image was also skeletonized and counted to determine total root length. Figure 2 illustrates how the phenes were extracted from the segmented image. Table 1 lists all extracted phenes and their descriptions. For further analysis, the image phenes were aggregated from all perspectives creating four more phenes. The Max. Max. Width is the maximum among the Maximum Widths of the plant root across all perspectives. The Min. Max. Width is the minimum among Maximum Widths of the plant root across all perspectives. The Eccentricity is defined as the ratio of Max. Max. Width to Min. Max. Width. The Max. Max. Width to Avg. Depth Ratio is the ratio of the Maximum Width from all perspectives to the average Depth from all perspectives. This phene is similar to Width-to-Depth Ratio but takes into account all the perspectives.

**Figure 2.**
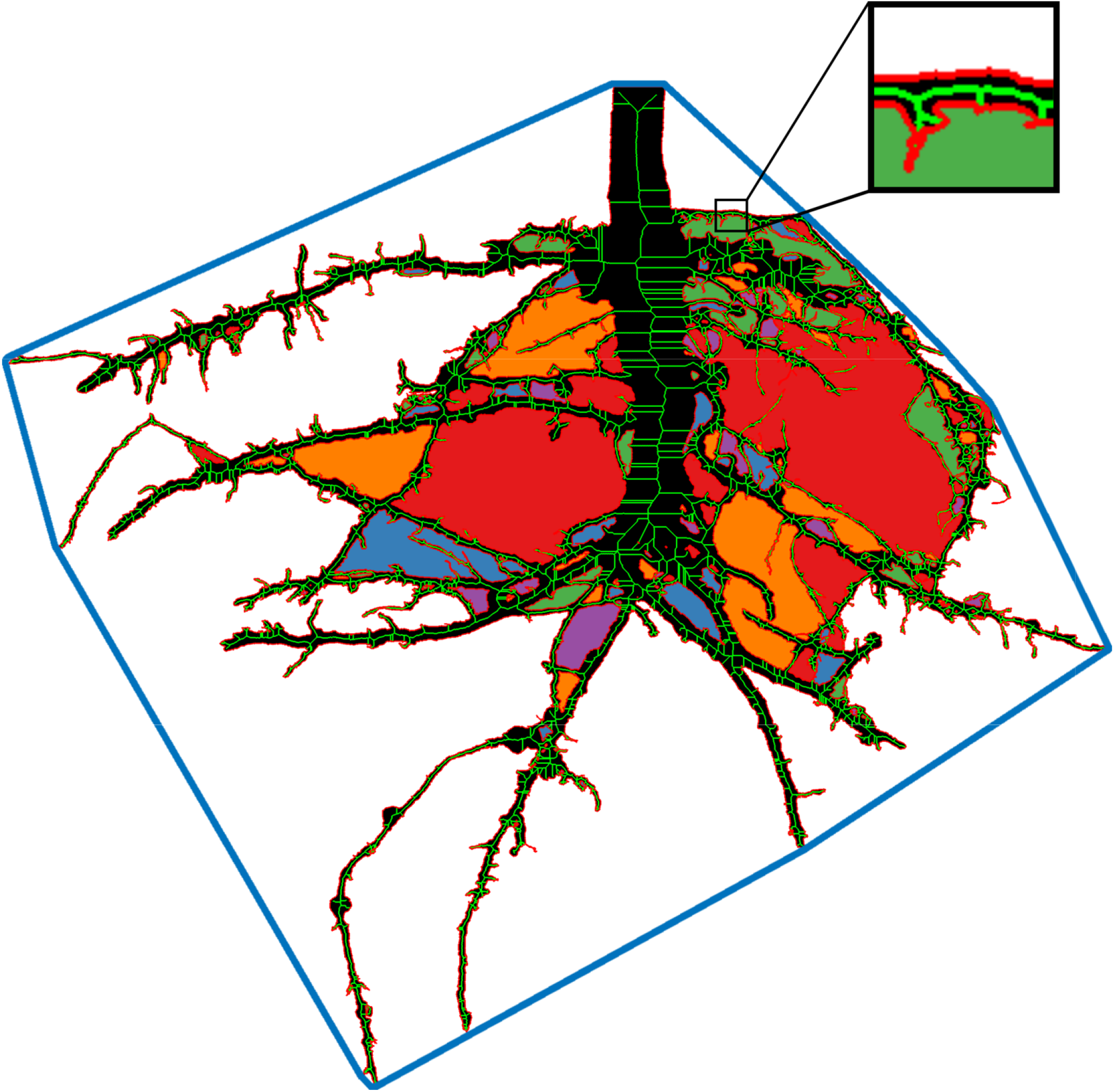
The EM-GMM algorithm generated a binary image with the root crown as a foreground object (black). Image analysis in MATLAB generated features such as the convex hull (blue line surrounding crown), the skeleton and length (green lines inside root segmented root crown), the perimeter of the root crown (red line along edge of root crown, see insert), and the number and size of disconnected components (multi-colored regions surrounded by the root crown edges).

**Table 1.**
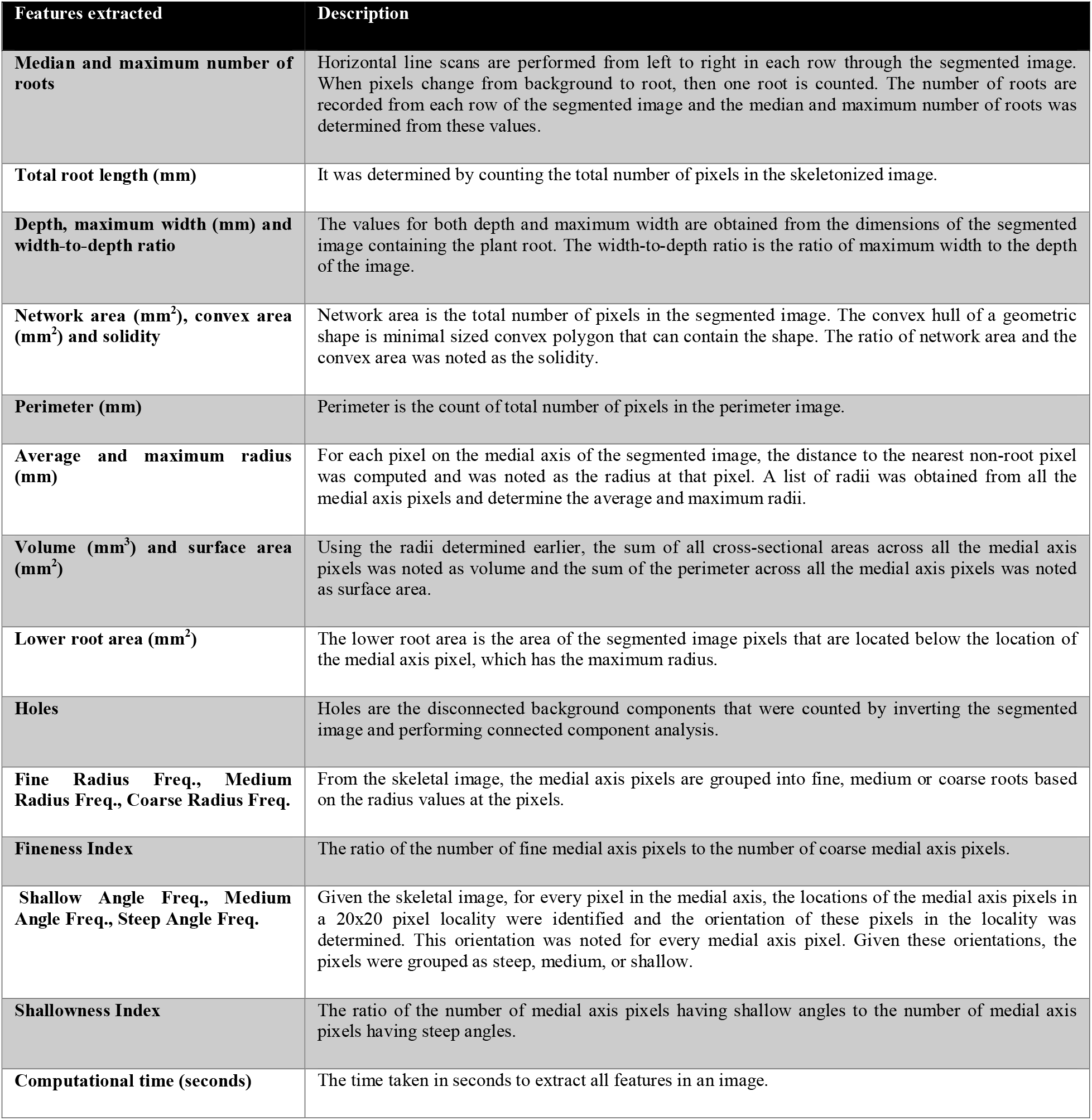
Features of root systems extracted from the segmented images with descriptions.

A total of 12 characteristics of the excavated root systems were manually measured or scored as described in Dhanapal et al. (under review), including Overall Complexity, Taproot prominence, Upper Primary Lateral Root Number, Upper Secondary Lateral Root Density, Upper Primary Lateral Root Angle Average, Upper Primary Lateral Angle Range, Lower Primary Lateral Root Number, Lower Secondary Lateral Root Density, Lower Primary Root Angle Average, and Lower Primary Lateral Angle Range, Total Number of Primary Lateral Roots, and Average Lateral Density. Additionally, Stem (1 cm above soil surface) and Tap Root Diameter (at 5 cm below uppermost primary lateral root), and Nodule Size (Average diameter of overall nodules compared to a scale of 1-5 mm) were measured and Nodule Density was scored (scale from 0 = no nodules to 5 = high Nodule Density).

### Statistical Analysis

Correlation analysis was performed among image phenes and manual phenes to explore the relation of automatically extracted phenes to manual phenes using data from individual root crowns (not averaged for plot). Correlation analysis was also performed for image phenes among different camera perspectives to check the validity of the phenes across perspectives. ANOVA was performed by taking the genotypes and perspectives as factors to test for significant effect sizes using data averaged within each plot. Using this analysis, the phenes that have large effect size for perspective factor may be identified as perspective-sensitive phenes. Finally, heritability calculations and MANOVA analysis for genotype partial effect size was performed on the extracted image phenes from all perspectives and manual phenes. Statistical analyses were performed in *R* (version 3.4). The functions *cor, mean* and *sd* were used for computing correlations, means of correlations and standard deviations (SD) of correlations respectively, where mean and standard deviation are given for data averaged within each plot. Similarly, *lm, anova* and *manova* were used to make linear models, perform ANOVA and MANOVA, respectively. In this study, linear models were used without any interactions.

Some phenes of roots may change as the perspective changes for the same plant root. For example, a root crown may have a smaller width when viewed from one angle compared to another angle (i.e., the root crown is approximately flat). In such a case, the extracted phenes from some perspectives may not correlate well with the manually acquired properties. To address this, the mean of the extracted phenes across all cameras was computed. Phenes were converted from pixel units to physical units before averaging the phenes, using camera sensor sizes and focal lengths.

To establish whether genotypes can be separated based on extracted phenotypes, ANOVA was performed on the phenotype data extracted from all five perspectives independently, the average phenotypes derived from the five perspectives, and the manual measures or scores. In total, 480 excavated roots were imaged for this study. For each of the 24 genotypes, roots of five plants were imaged from each plot (4 replications) and the plot averages of these five sub-samples were determined for each phene. Broad-sense heritability was calculated based on [2] as:

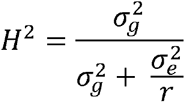

The variables 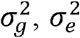, and *r* represent the variance of the genotype effect, variance of the environment effect, and the number of replicates (here, 4), respectively. The variances were obtained by fitting a mixed model including genotype as a random effect and replicate as a fixed effect using the *lme4* package.

Further, MANOVA was performed for each phene using all perspectives’ values as five response variables (Figure 8). The effect sizes in terms of partial eta squared was computed for each image phene using the following equation:

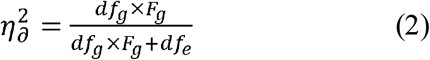

where df_g_ is the degrees of freedom for the genotype factor, F_g_ is the f-statistic for genotype factor and the df_e_ is the degree of freedom for residual error.

## Results

Twenty-five phenes were determined from image analysis of soybean root crowns excavated from the field (Table 1, Figure 2). The suite of image-based phenes covered various measures of size, length, radius distribution, branchiness, and angles. Phenes were also measured manually for comparison, as described in Dhanapal *et al*. (submitted). In total, 480 root crowns were imaged with five perspectives, yielding 2400 analyzed images.

Substantial variation in the population and field replicates existed for all measured phenes (Supp Table 1). Comparing means for each phene from each perspective and the averages across perspectives led to interesting patterns (Figure 3). As would be expected, the mean of Maximum Width was greatest for the center camera, while the other 4 perspectives were similar. The Depth was greatest for the farthest perspectives (Left 2 and Right 2), but smallest for the central perspective (V pattern). Lower Root Area means were relatively equal across perspectives (flat pattern). For many phenes, the mean of the central camera was most similar to the farther perspectives (W or M patterns).

**Figure 3.**
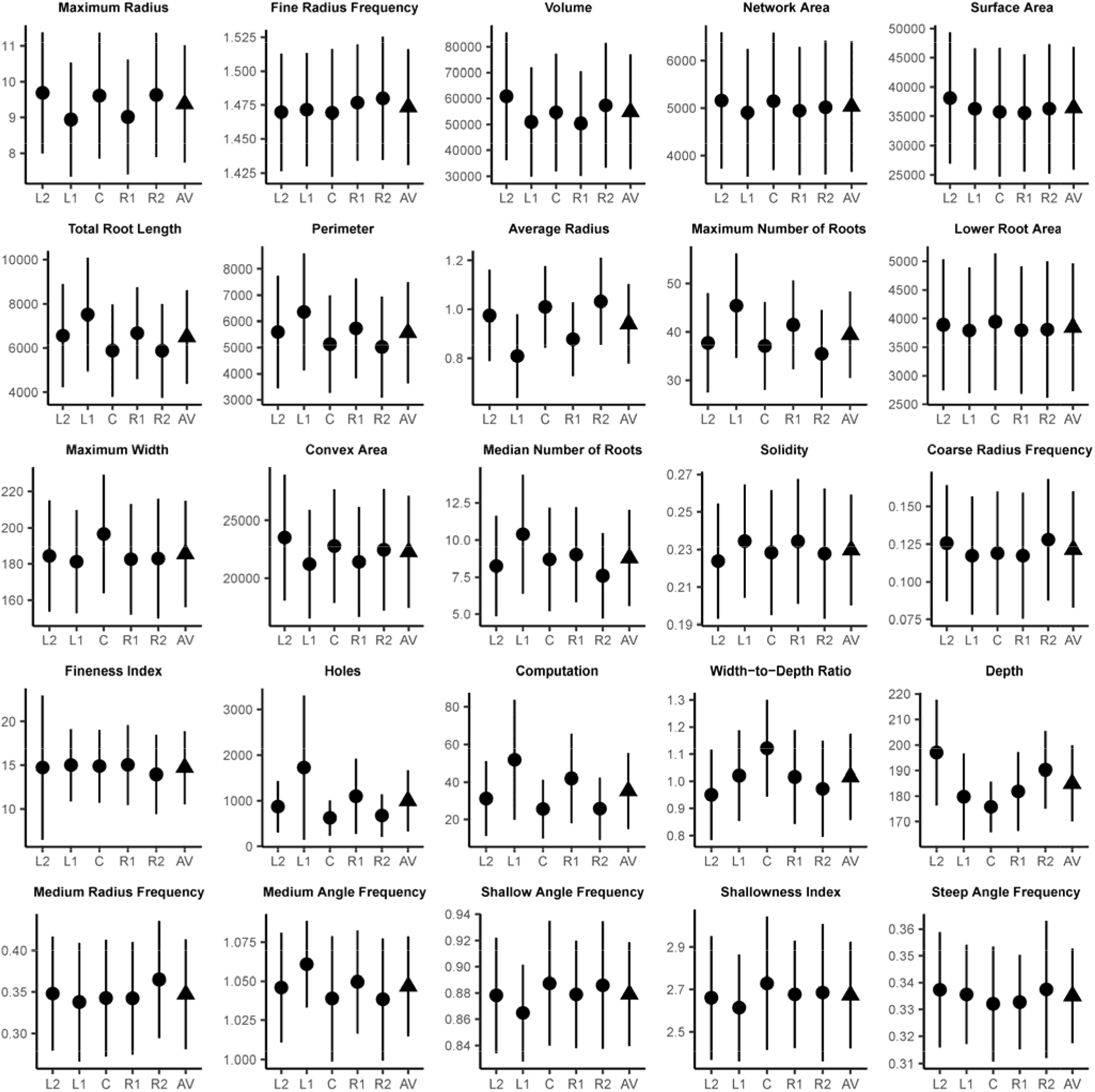
Means and standard deviations are given for the 25 image-based phenes, which are defined with respective physical units in Table 1. For each phene, mean +-SD is given for each camera perspective, Left 2 (L2), Left 1 (L1), Center (C), Right 1 (R1), and Right 2 (R2), all shown with black circles. Black triangles are the average (AV) and SD of each phene across the five perspectives.

### Correlation with the manually acquired properties

For each root, a total of 25 phenes were extracted from each image acquired by each camera. Correlation analysis was performed between image phenes and the manual phenes for each sample. The Pearson correlation coefficients and corresponding P-values were computed for each combination of the different image phenes and manual measures (Figure 4). The correlation analyses were performed (i) for each perspective (camera) separately, and (ii) based on the averages of the phenes obtained from the five perspectives. In addition, an average correlation coefficient was calculated based on the five single-perspective correlation coefficients. The maximum correlation for phenes averaged across the 5 perspectives was 0.67 for the image phene, Total Root Length, and the manual phene, Overall Complexity. The maximum correlation for individual perspectives (single camera) compared to manual measures was found to be 0.65, also for Total Root Length versus Overall Complexity. Similarly, the averaged image phenes across perspectives such as the Network Area, Perimeter, Lower Root Area and Computational Time correlated well with Overall Complexity with correlation coefficients of 0.65, 0.64, 0.61 and 0.63 respectively. The perspectives corresponding to the maximum correlations for each phene combination have no consistent pattern. Overall, there are correlations among image-based and manual phenes, yet it is not clear which should be most relevant as ground truth data.

**Figure 4.**
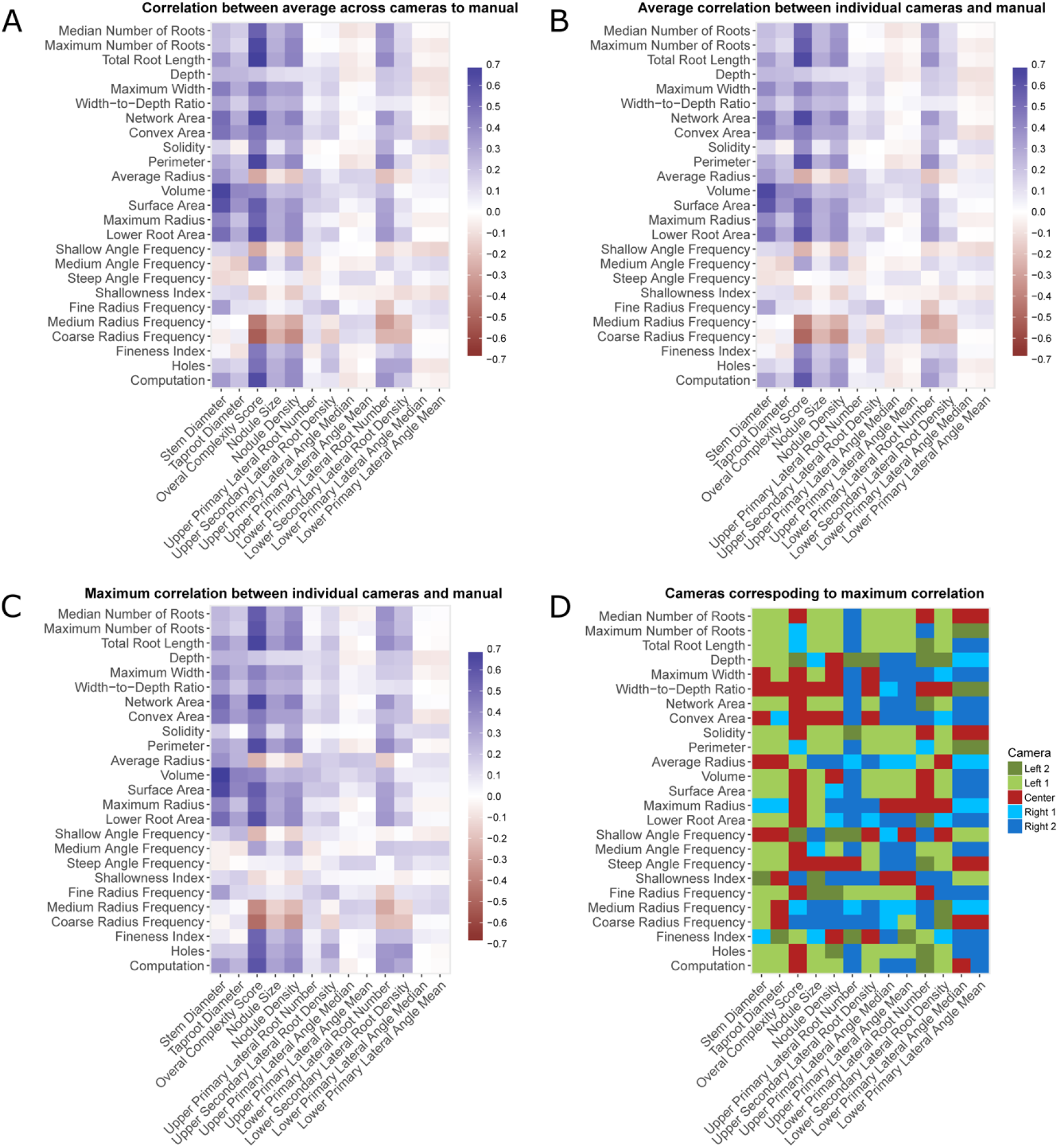
Pearson correlations calculated among digital and manual phenes. A. Correlations of phenes averaged across the five perspectives before correlational analysis with manual phenes. B. Averages of the five correlations from the five perspectives of each digital phene with manual phenes. C. Maximums of the five correlations from the five perspectives of each digital phene with manual phenes D. A map depicting which camera perspectives yielded the maximum correlations given in panel C.

In order to test whether averaging multiple perspectives was beneficial for predicting manual measurements, two operations were performed using the correlation tables described above. In Figure 5, the maximum correlations derived from individual perspectives were subtracted from the correlations of the average phene across 5 perspectives against manual phenes. In Figure 6, the averages of correlations derived from individual perspectives were subtracted from the correlations of the average phene across 5 perspectives against manual phenes. In both cases, positive numbers indicate the correlations from averaging phenes across 5 perspectives before correlational analysis with manual phenes performed better than the alternative. The averaged phenes did not correlate better than the maximum correlation values across all perspectives for most phene combinations (Figure 5), yet no single perspective had consistently greater correlations than averaging all perspectives (Figure 4). Averaging phenes across 5 perspectives before correlational analysis always performed better than the average correlations across multiple perspectives for phene combinations that are statistically significant (Figure 6). Since no single perspective consistently correlates better to manual phenes, and the best perspective is hard to predict *a priori*, averaging phenes across 5 perspectives before correlational analysis is recommended.

**Figure 5.**
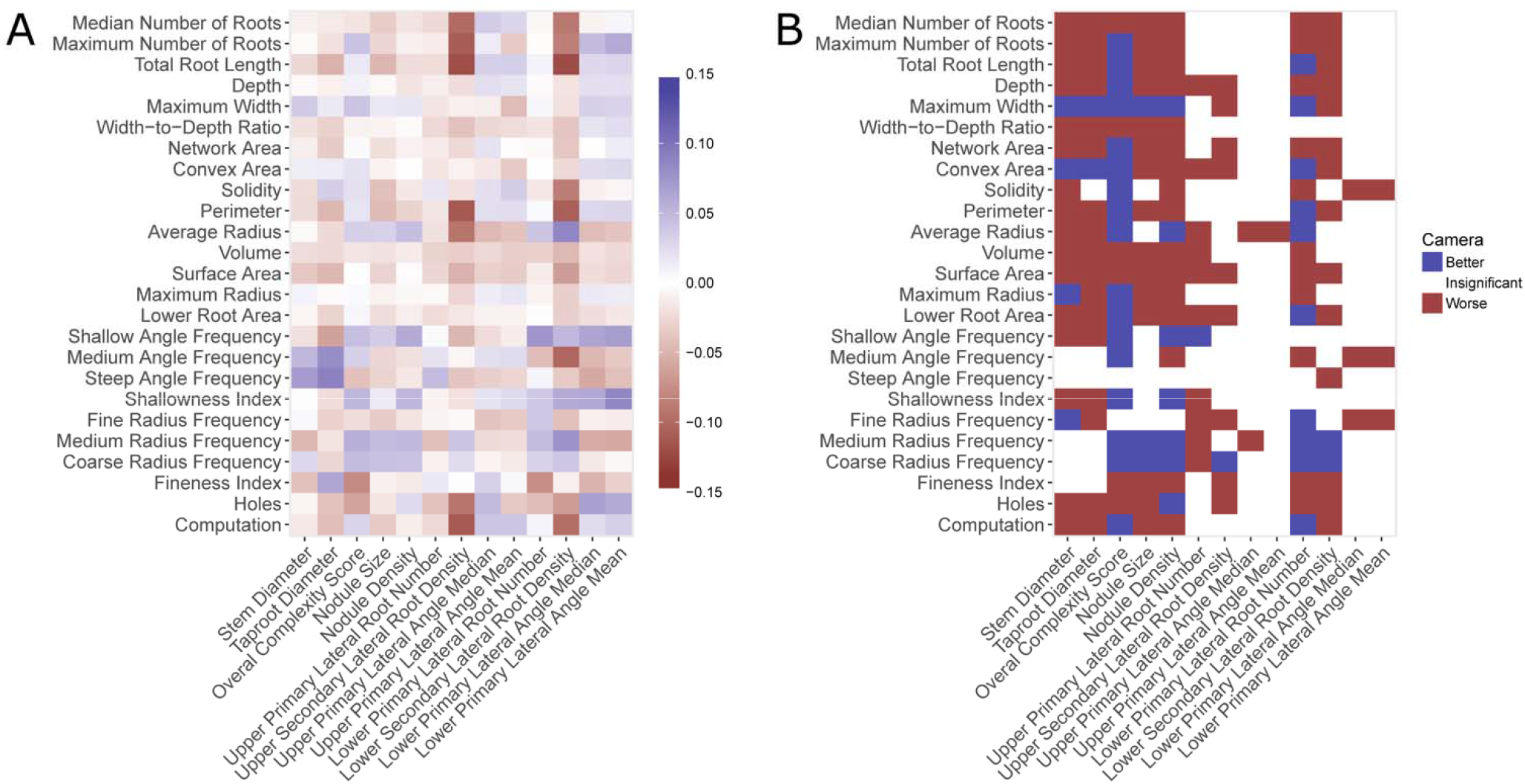
A. A heat map of the differences in correlations when the maximums of the five correlations from the five perspectives of each digital phene with manual phenes (Fig 3B) are subtracted from the correlations of phenes averaged across the five perspectives before correlational analysis with manual phenes (Fig 3A). B. Thresholded map demonstrating which differences of correlations indicate greater (better) correlations for averaging phenes across 5 perspectives before correlational analysis (only significant correlations shown).

**Figure 6.**
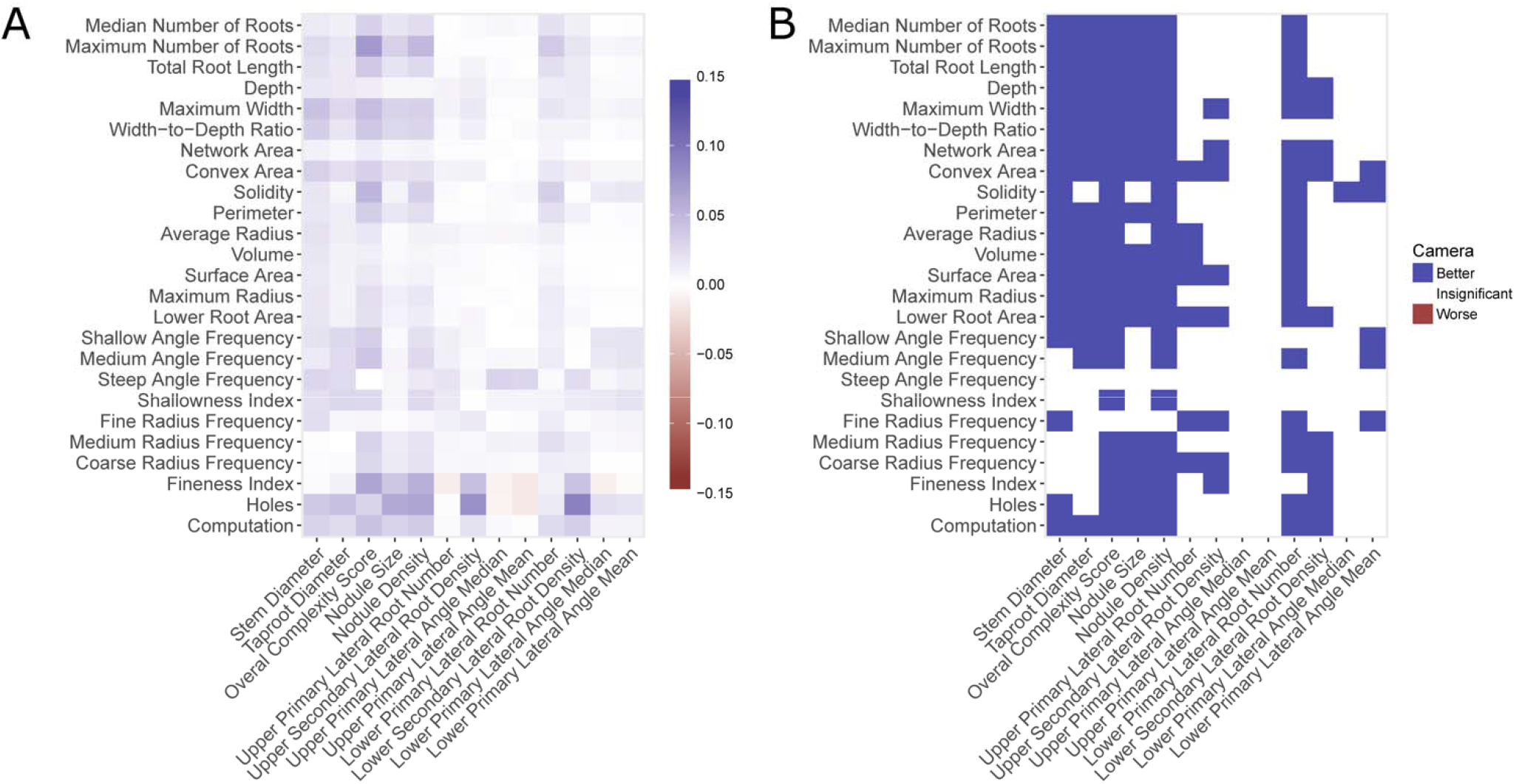
A. A heat map of the differences in correlations when the averages of the five correlations from the five perspectives of each digital phene with manual phenes (Fig 3C) are subtracted from the correlations of phenes averaged across the five perspectives before correlational analysis with manual phenes (Fig 3A). B. Thresholded map demonstrating which differences of correlations indicate greater (better) correlations for averaging features across 5 perspectives before correlational analysis (only significant correlations shown).

### Inter-perspective correlations

The correlations among the five perspectives for each image phene were computed (Figure 7). The maximum average inter-camera correlation was observed for network area, at 0.949±0.021 (mean±SD), followed by surface area (0.933±0.013) and volume (0.932±0.016). These substantial correlations indicate that even if the phene values are different from all the perspectives (slopes not equal to 1), the phene values change predictably across samples. The fineness index has the lowest average inter-camera correlation of 0.488±0.271, possibly a function of differential fine root occlusion among perspectives.

**Figure 7.**
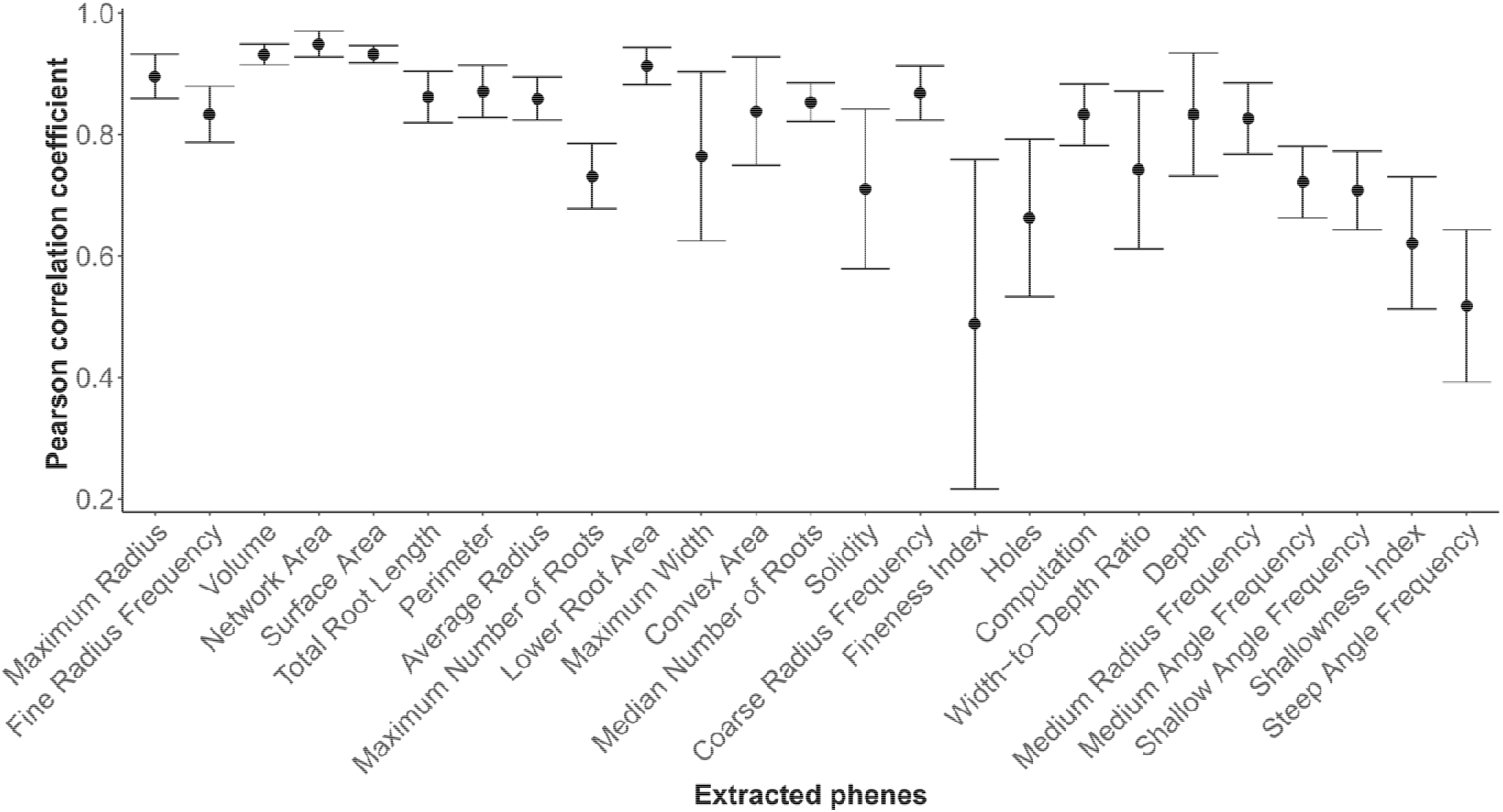
Averages and standard deviations for all pairwise correlations among the five perspectives for each image-based feature (n=10). Pearson correlations were calculated for all 25 pairs of perspectives before being averaged and the standard deviation calculated.

### Analysis of Variance Due to Perspective

Since the data was collected from five different perspectives for 24 soybean varieties, ANOVA was performed to partition variation created by the multiple factors in this experiment using eta-squared. The camera perspectives were also considered as a factor as the plant roots were oriented with the plane of maximum spread perpendicular to the central camera. Figure 8 shows the effect sizes of genotype, block in the field, and image perspective, as well as the residual of each image phene.

**Figure 8.**
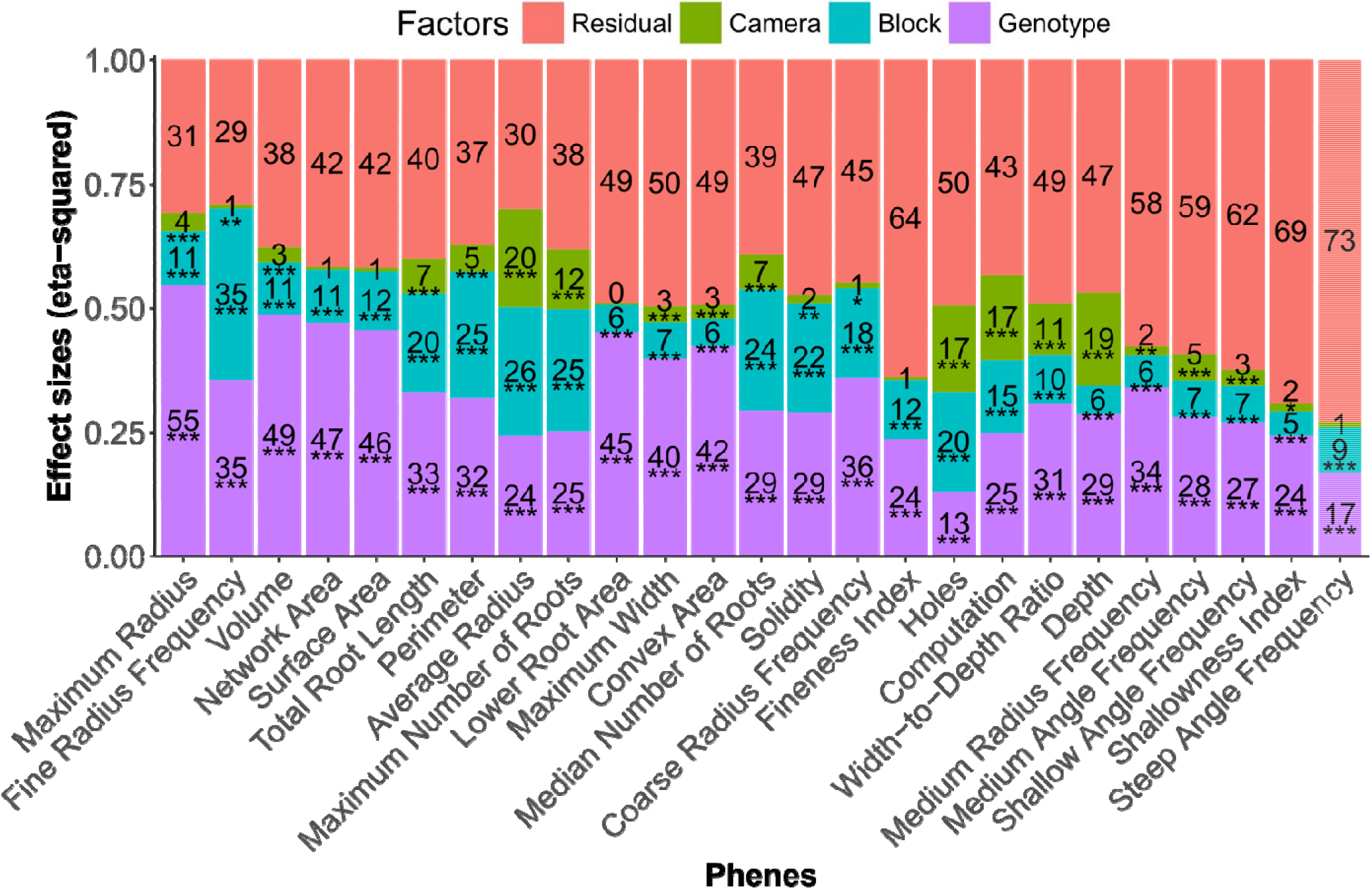
Effect sizes of all the factors for each phene extracted from the images. For each effect size, the percentage is shown that contributes to the total variation. Each effect size is marked with the significance of that factor (except for the residual error) based on p-values obtained. ‘*’ was given for p-value < 0.05, ‘**’ for p-value < 0.01 and ‘***’ for p-value < 0.001.

Genotype, block, and perspective effects were significant (P<0.001) for all phenes, except for the perspective effects for Volume, Surface Area, Lower Root Area, Fineness Index and Steep Angle Frequency (ns). The genotype effect size ranged from a low (13%) for Holes to a maximum (55%) for Maximum Radius. Thus, the Maximum Radius was particularly well suited for genotype separation. The phenes with the greatest effect sizes due to different perspectives were Average Radius, Holes, Computation and Depth. For the fineness index factor, the lower effect size associated with perspective agrees well with the large SD of the inter-camera correlations for the same phene in Figure 7.

### Heritabilities of image-based and manual phenes

Broad-sense heritability is an important metric for how suitable a new phene would be for breeding programs, assuming some phene states have beneficial effects on crop performance. The averaged phenes across perspectives generally had greater genotype heritability than phenes from individual perspectives (Figure 9). The greatest heritability for averaged image phenes was Maximum Radius with a heritability of 0.837 and the greatest heritability for manual phenes was Overall Complexity with heritability of 0.823. For comparing to the results of MANOVA, partial eta-squared was calculated, which is not directly comparable to heritability, but useful for looking at the differences among image-based phenes for separating genotypes using raw data from all 5 perspectives directly without averaging. The largest genotype effect size from MANOVA analysis of image phenes was 0.358 for Volume. The greatest heritability of derived phenes was observed fir Min. Max. Width with the heritability value of 0.693.

**Figure 9.**
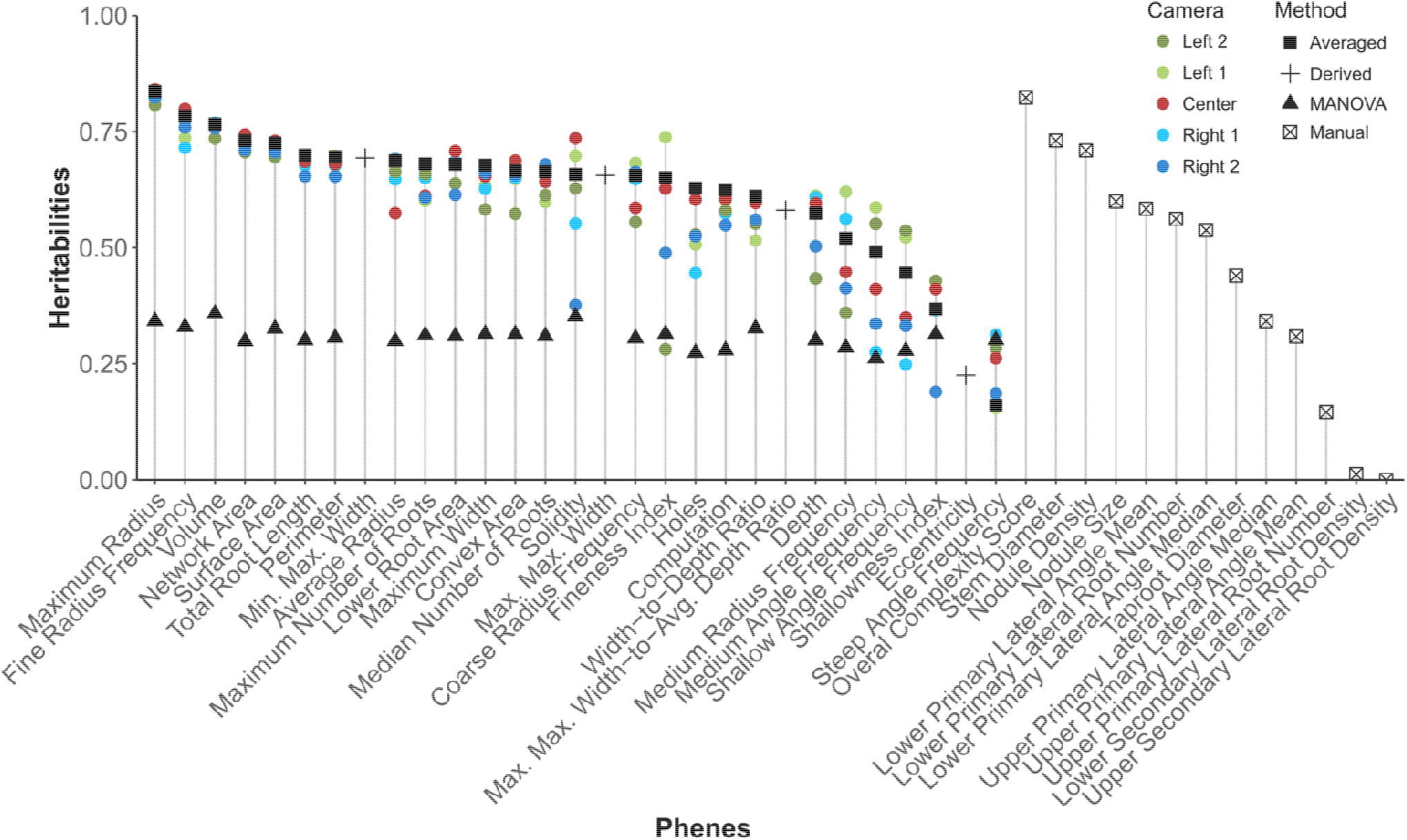
The broad-sense heritablities are plotted for individual camera perspectives (colored circles), using the average across perspectives (black squares), and with manual phenes draw win a black ‘x’ enclosed in a square. Effect-sizes for MANOVA (black triangles) using all five perspectives for each feature are displayed for comparison.

### Throughput of root crown imaging, segmentation, and phene extraction

The M-PIP system was tested for performance and compared to the existing systems such as *DIRT* (Bucksch et al. 2014) and *REST* (Colombi et al. 2015). The average imaging speeds of the M-PIP system was 1.66 root crowns min^-1^ whereas the imaging speed given for *REST* system was 1.5 root crowns min^-1^. Table 2 shows the segmentation time for various algorithms implemented for M-PIP, where the EM algorithm used here with GPU acceleration took 10 seconds to segment color images, while the DIRT implementation of Otsu’s thresholding took 22 seconds. On average the new feature extraction program, which was implemented in *MATLAB*, required 26 seconds for phene extraction on a PC with an Intel Xeon E3-1271 v3 processor (3.6 GHz) having 4 cores and 16 GB of RAM. An average of 860 seconds was required for phene extraction when the *DIRT* system was run in multi-threaded mode on PC running Ubuntu Linux with Intel Core i5-3440 processor (3.3 GHz) having 4 cores and 16 GB RAM.

**Table 2.** Average time required for the segmentation of all 2400 images across platforms. RVM and EM calculations were on a Intel Xeon E3-1271 v3 processor (3.6 GHz) having 4 cores and 16 GB of RAM, REST was reported in the original paper, and the DIRT implementation was run on an Intel Core i5-3440 processor (3.3 GHz) having 4 cores and 16 GB RAM.

| Segmentation Method | Average segmentation time (seconds) |
| --- | --- |
| RVM implementation in *MATLAB* | 90 |
| EM implementation in *MATLAB* | 50 |
| EM implementation in C++ with CPU acceleration | 25 |
| EM implementation in C++ with GPU acceleration | 10 |
| Image thresholding with trait extraction in REST system | 6 |
| DIRT implementation of image thresholding | 22 |

## Discussion

Image acquisition using the Multi-Perspective Imaging Platform (M-PIP) was faster than existing methods with regards to simultaneous imaging of the root crown from multiple perspectives, transferring the image data from each camera to the computer, and the time to place and replace root crowns. The setup box was operated by two persons, one sending commands to cameras to take pictures and the other replacing the plant roots after imaging. The two person team was able to keep up with the root crowns passed by a group of six (three teams with two people each) conducting manual measurements. Since, the setup box was designed to be invariant from external lighting conditions, the images had high contrast of root crown to background, leading to excellent segmentation outputs.

The EM algorithm used to segment the color images from the cameras provided more connected root pixels in the segmentation results compared to the segmentation by conventional thresholding algorithms on grayscale images. In this study, a blue-colored background was found to contrast best with roots and allowed the use of multi-dimensional color channel information to better segment roots from the background. This made the color channels in each image to store information for segmenting the images and hence lead to better results. The segmentation and phene extraction for the M-PIP was performed on a PC with an Intel Xeon E3-1271 v3 processor having 4 cores and 16 GB of RAM. The PC has a NVIDIA Quadro K600 GPU with 192 CUDA cores having Compute Capability 3.0 (a performance standard from NVIDIA). Optimization was achieved further by applying hardware acceleration techniques for CPU and GPU, making this program suitable for online segmentation of images. This direct segmentation of the images in the field enables immediate control of image quality and system performance, providing operator peace of mind and the ability to make adjustments if needed.

### Correlation with the manual phenes

The maximum correlation was increased after averaging phenes across the perspectives to 0.67 from the maximum correlation for individual perspectives of 0.65, for the feature combination of the image phene Total Root Length and the manual phene Overall Complexity. The overall complexity is scored manually based on apparent secondary and tertiary roots leading to more or less bushy structures. This means that a majority of the phenes extracted from the images are useful in predicting the complexity of the plant root automatically, and may allow complexity to be deconstructed to more basic properties. While the angle phenes extracted manually did not correlate significantly to the image phenes, the number of lower roots and the stem diameter correlated well. Since manual angles were measured on primary laterals whereas image-based angles were calculated for every pixel in the skeleton, including the taproot, secondary roots, and tertiary roots, the manual and image-based measures are fundamentally different. Additionally, angles may not be same as seen from various imaging perspectives and manual root angles were measured using a protractor in a perspective where the roots are spread widely, similar to the central camera. Additionally, occlusion of roots by other roots when viewed from particular perspectives may have contributed to the lack of significant correlations between manually measured angles and imaged based phene. The greatest negative correlations can be observed with the number of medium and coarse roots in a root crown with the overall complexity. This indicates that investment into large radii roots may decrease length and branching, leading to a decrease in overall complexity. The manual phenes Nodule Size and Nodule Density had minimal effect on overall complexity. Both manual and image-based measures may have relevance for agronomic performance, a focus of future work.

The camera perspectives of phenes that led to maximum correlations with the manual phenes were distributed with no obvious pattern (Fig. 3D), indicating that no single perspective consistently performed better than any other perspective. Although the average phenes across all perspectives did not perform as well as the maximum correlations selected from among the five perspectives (Fig. 4), the best performing perspective could not be predicted *a priori*. Hence, if the phenes from all the perspectives were averaged, an overall improvement in the correlation may be observed because the average of these phenes reduces the effect of the outliers that were present when the root crown was imaged from multiple perspectives. The average of the phenes from all the perspectives always out-performed the average correlations across all the perspectives. Therefore, averaging features across all five perspectives before analysis is recommended.

### Inter-camera correlations

The image phenes from various perspectives generally correlated well, except for a few phenes that significantly vary among perspectives. For example, finer roots easily can be blocked by coarse roots which may lead to a large variance in their numbers from different perspectives. This is shown in the Figure 7, where Fineness Index and holes had greater SD in correlations across all perspectives. Also, the orientation of the root either increases or decreases as a function of perspective. This may lead to misidentification of some shallow and medium angle roots as steep angle roots in some perspectives. While the SD for correlations of Maximum Width was reasonable because the phene varies with the change in perspective, a similar SD for correlations of Depth was curious. However, this may be attributed to the Depth of some roots exceeding the dimensions of the image in some perspectives which may be caused by non-precise alignment angles at which the cameras were mounted. Greater SD was also observed for phenes such as Convex Area and Solidity, which is reasonable as Maximum Width also had greater SD as perspective changed. The phenes with large inter-camera correlations and small SD such as network area, volume, and surface area etc. may be considered perspective insensitive. Hence, these phenes may be particularly well-suited and reliable when the root crowns are imaged from a single perspective. Greater SD for a phene correlation may imply that the perspectives may hold more information about the root crown such as the information needed for genotype separability.

### Factor separability of image-based phenes

Among image-based phenes, the absolute and relative effect sizes of the genotype, block, perspective, and residual error vary greatly (Figure 8). The phenes Average Radius, Holes, Computation and Depth have relatively larger perspective effect sizes. This was due to the larger width roots occluding the finer roots. As the perspectives changed the Average Radius and Holes changed and hence the Computational time also changed. Here, the Computational time may be approximately considered as proportional to the complexity of the root crown. When the perspective is changed the root may be recorded as smaller or larger size or having small or large number of Holes. If the root crown is large in size and the larger number of Holes, the Computational time increases. On the other hand, if the root crown is large in size and bushy such that the Holes are not visible, the Computational time decreases. This explains the similar perspective effect size of Computational time with Holes and the smaller genotype effect size of this phene, even though this phene correlates well with the manual phene Overall Complexity.

Frequency-based phenes had smaller effect sizes for the perspectives factor. This was attributed to the roots contain a majority of finer roots than coarse roots, thus counting of coarse roots from multiple perspectives may lead to similar values, thereby decreasing the effect size and significance. In case of Fine and Medium Radius Frequencies, the effects of occlusion, made the phenes less significant. In the Angle Frequencies, change in the perspectives also changes the angle of the roots. This substantial dependency on the change in perspective coupled with the problems due to occlusion resulted in a smaller effect size of Frequency phenes for the perspective factor.

The genotype factor generally had a larger effect than perspective and block. Interestingly, the phenes that were most susceptible to change in perspective were not well-suited to separate genotypes. On the other hand, many of the phenes that were susceptible to perspective yielded greater partial effect sizes when performing MANOVA using phenes from all the perspectives to test for genotype separability (Figure 9).

### Heritabilities of image-based and manual phenes

Heritability and MANOVA results revealed that the image phene maximum radius had a greater heritability than the manually scored Overall Complexity (Figure 9). For genotype separation, the averaged phenes from all camera perspectives generally perform better than the phenes from individual perspectives. This shows that imaging roots from multiple perspectives may provide information which may otherwise be lost or limited due to occlusion of roots and changes in angle based on perspective. To examine the influence of multiple perspectives for genotype separability, MANOVA was performed by taking the phene values from across all the perspectives. The Volume and Solidity phenes had the greatest MANOVA partial effect-sizes compared to the remaining phenes. Interestingly, the phenes having larger effect-sizes for the perspective factor did not have higher MANOVA partial effect size (from Figure 9). This indicates that effect-sizes in perspective factors for these phenes did not contain information for genotype separability. On the other hand, the phenes Volume and Solidity have larger effect-sizes in MANOVA analysis than using phenes from single perspectives for heritability. This shows that including each perspective in the analysis for these phenes can increase genotype separability using MANOVA. The phenes for the Angle Frequencies had smaller heritabilities and MANOVA partial effect sizes. This may be attributed to the change in orientation of the roots as the perspective changed. Also, both Volume and Solidity are susceptible to occlusion, likely contributing to smaller partial effect-sizes in MANOVA analysis. In the aggregated features combined from the phenes from all the perspectives, the phene Min. Max. Width had a greater heritability than the Maximum Width, and that of the phene Max. Max. Width had smaller heritability. While these aggregated phenes have very similar heritabilities, more studies are needed to conclude that these aggregated phenes can be used reliably in root crown phenotyping.

The similarity of heritabilities of image phenes and those of manual phenes indicates that the image phenes can be used reliably. The image phenes can be investigated further to better understand their role or importance for plant growth and performance in different environments. The phenes with the greatest heritabilities may be the most useful for breeding, but more needs to be known about their relation to crop performance.

## Conclusions

The Multi-Perspective Imaging Platform described here is capable of imaging hundreds of root crowns a day from five perspectives. A reliable segmentation algorithm which can efficiently retrieve the root structure from color images was implemented. The segmentation program utilized hardware acceleration leveraging CPU and GPU resources present on the PC to achieve faster segmentation rates without compromising on the quality of the segmentation process. A variety of root phenes were extracted, some are based on the size and appearance of the plant root, while other phenes were intended to extract hidden structures or additional information about the roots. Image acquisition in the field was faster than documented for the REST platform, and image analysis was faster than DIRT but slower than REST (which used greyscale thresholding). Finally, using phenes across perspectives allowed both greater correlations to manual phenes and greater heritabilities relative to only using individual perspectives.

Future improvements in image phene extraction include implementation of new phenes such as root angles from each root class and novel ways of merging phenes from multiple perspectives instead of using the mean across all camera perspectives. Such advances may result in greater precision when used for genetic mapping and thus more reliable genetic markers for breeding purposes. Furthermore, more integration of hardware and software specifically for root crown phenotyping has potential to greatly increase throughput and reliability.

The M-PIP facilitates rapid, quantitative assessment of root phenes and can be used in physiological experiments to link root phenotypes to measures of crop performance such as grain yield, nutrient content, and shoot mass. Likewise, the phenes can be mapped to genetic regions using QTL analysis and GWAS to enable marker assisted breeding for root phenes. As such, the M-PIP system can facilitate the inclusion of specific root phenes as targets for breeding programs to aid in the development of more productive, stress tolerant crops and reduce environmental impact due to nutrient loss from agricultural ecosystems.

## List of Abbreviations

ANOVA: – Analysis of Variance
CHDK: – Canon Hacker Development Kit
EM: – Expectation Maximization
M-PIP: – Multi-Perspective Imaging Platform
MANOVA: – Multi-variate Analysis of Variance
PTP: – Picture Transfer Protocol
RCBD: – Randomized Complete Block Design
RSA: – Root System Architecture

## Declarations

### Availability of data and materials

Raw images and/or segmented masks are available upon request. Software code is available on github (DOI: 10.5281/zenodo.1213805 | website: https://github.com/GatorSense/MPIP).

### Restrictions or Required Licenses

No restrictions on this research are known under local or national laws.

### Authors’ contributions

AS wrote the software, imaged root crowns, conducted data analysis, and wrote the manuscript. LY provided input to software, built the current imaging platform, imaged root crowns, gave input to data analysis, and wrote the manuscript. HA managed the field experiments, helped build the platform, measured manual phenes, helped analyze data, and helped write the manuscript. FF and AZ conceived of the project, directed the design of the imaging platform, directed software design, gave input to data analysis, and wrote the manuscript.

## Acknowledgements

We appreciated the help of Arun Dhanapal and Xiaoxiao Du in collecting samples and providing the manual measurements.

## References

Balestri E, de Battisti D, Vallerini F, Lardicci C. First evidence of root morphological and architectural variations in young Posidonia oceanica plants colonizing different substrate typologies. Estuarine, Coastal and Shelf Science. 2015;154:205–213.

Basu P, Pal A, Lynch JP, Brown KM. A novel image-analysis technique for kinematic study of growth and curvature. Plant Physiology. 2007;145:305–316.

Beentje H. The Kew Plant Glossary: an illustrated dictionary of plant terms. Royal Botanic Gardens, Kew, Richmond, UK. 2010.

Bilmes JA. A gentle tutorial of the EM algorithm and its application to parameter estimation for Gaussian mixture and hidden Markov models. International Computer Science Institute. 1998.

Bishop CM. Pattern Recognition and Machine Learning (Information Science and Statistics). Springer-Verlag New York, Inc. 2006.

Bucksch A, Burridge J, York LM, Das A, Nord E, Weitz JS, Lynch JP. Image-Based High-Throughput Field Phenotyping of Crop Roots. Plant Physiology. 2014;166:470–486.

Burridge J, Jochua CN, Bucksch A, Lynch JP. Legume shovelomics: High—Throughput phenotyping of common bean (Phaseolus vulgaris L.) and cowpea (Vigna unguiculata subsp, unguiculata) root architecture in the field. Field Crops Research. 2016;192:21–32.

Burridge JD, Schneider HM, Huynh BL, Roberts PA, Bucksch A, Lynch JP. Genome-wide association mapping and agronomic impact of cowpea root architecture. Theoretical and Applied Genetics. 2016b; 130:1–13.

CHDK CHDK Home. http://chdk.wikia.com/wiki/CHDK.

Chen YH, Zhang Q, Li BG, Zhang BG. Characterizing Wheat Root Branching Using a Markov Chain Approach. Second International Symposium on Plant Growth Modeling and Applications, Beijing, 2006. 70–73.

Clark RT, MacCurdy RB, Jung JK, Shaff JE, McCouch SR, Aneshansley DJ, Kochian LV. Threedimensional root phenotyping with a novel imaging and software platform. Plant Physiology. 2011. 156:455–465.

Colombi T, Kirchgessner N, Le Marié CA, York LM, Lynch JP, Hund A. Next generation shovelomics: set up a tent and REST. Plant and Soil. 2015;388 (1):1–20.

Dempster AP, Laird NM, Rubin DB. Maximum Likelihood from Incomplete Data via the EM Algorithm. Journal of the Royal Statistical Society Series B (Methodological). 2015;39:1–38

Dupuy L, Fourcaud T, Stokes A, Danjon F. A density-based approach for the modelling of root architecture: application to Maritime pine (Pinus pinaster Ait.) root systems. Journal of Theorhetical Biology. 2005;236:323–334.

Falconer D, Mackay T. Introduction to quantitative genetics. Essex, UK: Longman Group Ltd. 1996.

Fang Y, Wenyong W, Shaochun Z, Ye T, Linan Y. Modeling and Research on Growth of Virtual Plant Wheat Roots. International Forum on Information Technology and Applications, Chengdu. 2009. 264–267.

Fehr WR, Caviness CE. Stages of soybean development. Special Report 80. Iowa State University Cooperative Extension Service. Ames, IA. 1977.

Fenta B, Beebe S, Kunert K, Burridge J, Barlow K, Lynch J, Foyer C. Field Phenotyping of Soybean Roots for Drought Stress Tolerance. Agronomy. 2014;4:418–435.

Grafton RQ, Williams J, Jiang Q. Food and water gaps to 2050: preliminary results from the global food and water system (GFWS) platform. Food Security. 2015;7:209–220.

Huang Q, Jain AK, Stockman GC, Smucker AJM Automatic image analysis of plant root structures. In: Proceedings., 11th IAPR International Conference on Pattern Recognition. Vol.II. Conference B: Pattern Recognition Methodology and Systems. 1992. 569–572.

Iyer-Pascuzzi AS, Symonova O, Mileyko Y, Hao Y, Belcher H, Harer J, Weitz JS, Benfey PN. Imaging and analysis platform for automatic phenotyping and trait ranking of plant root systems. Plant Physiology. 2010;152:1148–1157.

Janusch I, Kropatsch WG, Busch W. Topological Image Analysis and (Normalised) Representations for Plant Phenotyping. 16th International Symposium on Symbolic and Numeric Algorithms for Scientific Computing, Timisoar. 2014. 579–586.

Jia Y, Su Z, Sun H. Research on the model construction of soybean root system based on L-system. World Automation Congress. 2010. 195–199.

Kirchgessner N, Spies H, Scharr H, Schurr U Root growth analysis in physiological coordinates. Proceedings 11th International Conference on Image Analysis and Processing. 2001. 589–594.

Kun B, Feifei H, Xin Z, Cheng W. Image Collection Filed of Plant Roots Based on Hypoid Mirror. Fourth International Conference on Intelligent Computation Technology and Automation, Shenzhen. 2011. 563–567.

Lobet G, Pagès L, Draye X. A Novel Image-Analysis Toolbox Enabling Quantitative Analysis of Root System Architecture. Plant Physiology. 2011;157:29–39.

Lynch JP. Root architecture and plant productivity. Plant Physiology. 1995;109:7–13

Mairhofer S, Zappala S, Tracy SR, Sturrock C, Bennett M, Mooney SJ, Pridmore T. RooTrak: Automated Recovery of Three-Dimensional Plant Root Architecture in Soil from X-Ray Microcomputed Tomography Images Using Visual Tracking. Plant Physiology. 2012;158:561–569.

Pagès L, Bécel C, Boukcim H, Moreau D, Nguyen C, Voisin A-S. Calibration and evaluation of ArchiSimple, a simple model of root system architecture. Ecological Modelling. 2014;290:76–84.

Pantalone VR, Rebetzke GJ, Burton JW, Carter TE. Phenotypic Evaluation of Root Traits in Soybean and Applicability to Plant Breeding. Crop Science. 1996;36:456–459.

Pound MP, French AP, Atkinson JA, Wells DM, Bennett MJ, Pridmore T. RootNav: Navigating Images of Complex Root Architectures. Plant Physiology. 2013;162:1802–1814.

Saengwilai P, Tian X, Lynch JP. Low crown root number enhances nitrogen acquisition from low nitrogen soils in maize (Zea mays L.). Plant Physiology. 2014;166:581–589.

Topp CN, Iyer-Pascuzzi AS, Anderson JT, Lee CR, Zurek PR, Symonova O, Zheng Y, Bucksch A, Mileyko Y, Galkovskyi T, Moore BT, Harer J, Edelsbrunner H, Mitchell-Olds T, Weitz JS, Benfey PN. 3D phenotyping and quantitative trait locus mapping identify core regions of the rice genome controlling root architecture. PNAS. 2013;110:1695–1704.

Trachsel S, Kaeppler SM, Brown KM, Lynch J Shovelomics: high throughput phenotyping of maize (Zea mays L.) root architecture in the field. Plant and Soil. 2011;341:75–87.

Trachsel S, Kaeppler SM, Brown KM, Lynch JP. Maize root growth angles become steeper under low N conditions. Field Crops Research. 2013;140:18–31.

USDA FAS Oilseeds: World Markets and Trade. https://www.fas.usda.gov/data/oilseeds-world-markets-and-trade.2018.

Ying Z, Gu S, Edelsbrunner H, Tomasi C, Benfey P Detailed reconstruction of 3D plant root shape. International Conference on Computer Vision. 2011. 2026–2033.

York LM, Galindo-Castaneda T, Schussler JR, Lynch JP. Evolution of US maize (Zea mays L.) root architectural and anatomical phenes over the past 100 years corresponds to increased tolerance of nitrogen stress. Journal of Experimental Botany. 2015;66:2347–2358.

York LM, Lynch JP. Intensive field phenotyping of maize (Zea mays L.) root crowns identifies phenes and phene integration associated with plant growth and nitrogen acquisition. Journal of Experimental Botany. 2015;66:5493–5505.

York LM, Nord EA, Lynch JP. Integration of root phenes for soil resource acquisition. Frontiers in Plant Science. 2013;4:1–15.

Yugan C, Xuecheng Z Plant root image processing and analysis based on 2D scanner. Fifth International Conference on Bio-Inspired Computing: Theories and Application. 2010. 1216–1220.

Zhou X, Cao X, Zhang C, Yan H, Li Y, Luo X. A method of 3D nondestructive detection for plant root in situ based on CBCT imaging. 7th International Conference on Biomedical Engineering and Informatics. 2014. 110–115.

